# Predicting West Nile Virus risk across Europe for the current and future conditions

**DOI:** 10.1101/2025.03.04.640552

**Authors:** A. J. Withers, S. Croft, R. Budgey, D. Warren, N. Johnson

**Author notes:** **corresponding authors:** /. **data availability statement:** All data underlying this work are held in publicly accessible repositories and can be downloaded for use in research under licence. Outputs can be made available under license by contacting the corresponding author. **funding statement:** This work is funded by EXSE0566: Development of improved tools and approaches for control of zoonotic arthropod-borne diseases of animals and their vectors Funded by the Department for Environment, Food and Rural Affairs and the Scottish and Welsh Governments and was co-funded by the European Union’s Horizon Europe Project 101136346 EUPAHW. **ethics approval statement:** n/a. **patient consent statement:** n/a. **permission to reproduce material from other sources:** n/a. **clinical trial registration:** n/a.

## Abstract

Vector-borne diseases have significant impacts on animal and human health globally, and these impacts are likely to increase in the future due to environmental and climate change. Understanding where to target surveillance and control measures to mitigate the impacts of emerging vector-borne diseases can be challenging when pathogens or disease is absent. In this study, we utilise a species distribution modelling approach previously applied to the UK to predict areas at higher risk of mosquito-borne disease across Europe, using West Nile Virus (WNV) as a case study. WNV is an *Orthoflavivirus* that is naturally transmitted between *Culex* mosquitoes and a range of avian species. However, it can spread to hosts such as humans and horses where it has the potential to lead to severe illness and mortality. Suitability predictions for *Culex* (*Cx. pipiens* and *Cx. modestus*) and avian hosts (mainly Passerine species) are made across Europe to determine potential risk of WNV circulation and establishment. These maps are then combined with information on human and horse density to determine risk to human and equine health. The resulting risk maps reveal that across Europe, there are areas of higher and lower risk that are predominantly driven by vector suitability as avian hosts are widespread. These predictions are projected into the future in 2100 using best- and worst-case Shared Socioeconomic Pathways (SSP1 and SSP5 respectively) to determine how risk may change over time, revealing that some areas see an increase in suitability for both vectors and hosts leading to higher risk (e.g., central England and northern Belgium) whilst other areas see a decline in suitability and consequently lower WNV risk (e.g., northern Italy and western Germany). Overall, this work will improve understanding of mosquito-borne disease risk in changing environments and demonstrates how species distribution modelling can be used to aid contingency preparedness by highlighting areas at higher risk of emerging disease.

## Introduction

Vector-borne diseases are an increasing threat to human and animal health globally as range shifts are occurring, with many factors such as climate change, increased movements, socio-economic factors and land-use change all contributing to this increased risk (World Health Organisation 2020, Endo & Amarasekare 2022, Rocklöv & Dubrow 2020, Noll et al. 2023). Understanding areas at greater risk of vector-borne diseases can focus surveillance and mitigation measures, reducing the impacts on human and animal health as well as lowering the economic cost (World Health Organisation 2020, Fabuary 2015). However, mitigating against potential risk, in regions where vectors and/or pathogens are not currently present, is challenging.

Understanding the risk of vector-borne disease requires an understanding of both host and vector distributions, and one way that this can be achieved is through species distribution modelling. Species distribution modelling has been used for a wide range of taxa, including mammals, birds and insects. It can predict the suitability of a landscape for a species providing an indication of its potential range and abundance, which can be particularly beneficial for estimating the potential range of a species or disease that is not yet present in an area, as well as improving understanding of native species when limited information in available (Amdouni et al. 2022, Baici & Bowman 2023, Chapman et al. 2019, Croft et al. 2017, Early et al. 2022, Elith et al. 2010, Velu et al. 2023). By providing estimates of where is most suitable for vectors the potential risk of diseases to human or animal health can be estimated, for example species distribution modelling has been extensively used for *Aedes* mosquito species and various diseases in different regions (Khormi & Kuma 2014, Kraemer et al. 2019, Oliveira et al. 2023, Richman et al. 2018, Beebe et al. 2009).

There are two types of species distribution models used for modelling insect distributions, these are correlative and mechanistic (Koch 2021). Correlative models determine suitability based on occurrence data and a set of environmental variables, and then this suitability model is used to predict suitability across a wider study area (Koch 2021, Elith et al. 2010, Hijmans & Elith 2013, Valavi et al. 2021). Conversely, mechanistic models do not require occurrence data as they are based on *a priori* knowledge of a species tolerance to environmental stressors, survival constraints and seasonal population growth to predict the species response to climatic variables, and as such are most suitable for species with strong climatic dependencies such as plants and insects (Koch 2021, Sutherst & Maywald 1985, Kriticos et al. 2015). For both correlative and mechanistic models there a number of different modelling approaches available, and selecting the most appropriate can be challenging as each have their own merits depending on the species and data available (Koch 2021, European Centre for Disease Prevention and Control 2019, Simons et al. 2019, Furlong et al. 2022, Gorris et al. 2021).

Previously, we have approached predicting WNV risk across the UK through using a combination of mechanistic and correlative species distribution modelling to determine risk to human and equine health (Withers et al. 2024). West Nile Virus (WNV) is an orthoflavivirus that is naturally transmitted by *Culex* mosquitoes (*Culex pipiens* and *Culex modestus*) biting birds. However, it can occasionally infect non-target hosts if bitten by infected mosquitoes, such as humans and equids. In humans, infection is usually asymptomatic however on rare occasions infection can be mildly symptomatic (e.g., headache, rash), or severely symptomatic in around 1 in 150 patients (e.g., acute aseptic meningitis) leading to illness or death (Pradier et al. 2012). Similarly, in horses WNV is usually asymptomatic but rarely it will lead to severe illness if it becomes neuroinvasive (Ostlund et al. 2000). As well as devastating impacts to human and animal health, WNV can have significant impacts on the economy, due to the high costs associated with treating symptomatic humans and mitigating the impacts to the equestrian sector, with predictions of the economic impact in Belgium of over €17million for the vaccination of horses in at-risk areas (Humblet et al. 2016). In recent years there have been WNV outbreaks in Europe where it has not been present historically, including France and Germany (Vogels et al. 2017, Constant et al. 2022, Bessell et al. 2016). Therefore, understanding the risk of WNV across Europe is important for contingency preparedness and policy decision making to mitigate and minimise the potential impact of this disease as it expands its range.

Here, our approach is expanded to predict risk across Europe, both in the current climate and in the future using two shared socio-economic pathways (SSP) to represent the best case (SSP1, climate protection measures and development compatible with 2°C target) and worst case (SSP5, no climate protection measures to work towards achieving the 2°C target) scenario in 2100. This can help determine locations to target surveillance to ensure early detection of WNV introduction to a pathogen-free area.

## Methods

Previous work used a combination of mechanistic and correlative species distribution modelling to produce models of habitat suitability for vectors and hosts across the UK (Withers et al. 2024). This was then combined to produce overall risk maps based on vector-host suitability and estimates of human and horse density added to give an overall indication of risk. This process of combining suitability predictions for vectors and hosts was replicated here using the models developed in Withers et al. (2024), where a full methodology on their development can be found. Here, these models are projected across a wider area to cover much of Europe for the current climate, as well as considering how this risk could change by 2100 under a best-case Shared Socioeconomic Pathway (SSP1) and worst-case (SSP5) future climate scenario.

In brief, global presence data for vectors *(Culex pipiens* and *Culex modestus)* and avian hosts (common raven *(Corvus corax),* rook *(Corvus frugilegus)*, carrion crow *(Corvus corone),* house sparrow *(Passer domesticus),* blackbird *(Turdus merula),* barn swallow *(Hereunto rustica)),* were obtained, cleaned and finally filtered based on environmental suitability (NBN Atlas 2022, GBIF 2022, 2024, 2024a, 2024b, 2024c, Culverwell & Vapalahti 2023). Random background points were generated in R for both vectors and hosts (dismo::*randomPoints,* v1.3-14, (Hijmans et al. 2023)). For vectors, a 100km buffer around presence points was determined for the random background points to be allocated in (Withers et al. 2024). For hosts, random background points were allocated within a convex polygon around presence points (Withers et al. 2024). Overall, for *Cx. pipiens*, this yielded 1319 presence and 3064 background points for model training and 253 presence and 1531 background points reserved for testing. For *Cx. modestus,* 27 presence and 72 background points for training and 21 presence and 70 background points for testing were used. For hosts, individual species data were combined for modelling into non-/corvid representative groups. For corvids there were 33,132 (22,994 and 10,138 in the training and testing datasets respectively) presences and 32,002 (19,476 and 12,526 in the training and testing datasets respectively) background points. For non-corvids there were 43,085 (25,944 and 17,141 in the training and testing datasets respectively) presences and 42,273 (22,108 and 20,165 in the training and testing datasets respectively) background points.

For vectors, a mechanistic CLIMEX model was produced as well as six correlative models (v. 4, (Sutherst & Maywald 1985, Kriticos et al. 2015)). These correlative models were Random Forest (RF, flexsdm::*tune_raf, v1.3.4, Velazco et al. 2022*), Maxent (MAX, flexsdm::*tune_max, v1.3.4, Velazco et al. 2022*), Generalized Boosted Regression (GBM, flexsdm::*tune_gbm, v1.3.4, Velazco et al. 2022*), Neural Network (NET, flexsdm::*tune_net, v1.3.4, Velazco et al. 2022*), Support Vector Machine (SVM, flexsdm::*tune_svm, v1.3.4, Velazco et al. 2022*) and an Ensemble model based on all five individual models (flexsdm::fit_ensemble*, v1.3.4, Velazco et al. 2022).* Model selection of the correlative models was based on the best performing model across several performance measures and was validated by comparing to the CLIMEX model. These performance measures were True positive rate (TPR), True negative rate (TNR), Sorensen, Jaccard, F-measure of presence-background (FPB), Omission rate (OR), True skill statistic (TSS), Area under curve (AUC), Boyce index and Inverse mean absolute error (IMAE) (Velazco et al. 2022, Valavi et al. 2022, Valavi et al. 2021, Hijmans & Elith 2013, Roberts et al. 2017).

Four correlative models were fitted for avian hosts, these were Random Forest (RF, flexsdm::*tune_raf,* v1.3.4, Velazco et al. 2022), Generalized Boosted Regression (GBM, flexsdm::*tune_gbm,* v1.3.4, Velazco et al. 2022), Neural Network (NET, flexsdm::*tune_net, v1.3.4,* Velazco et al. 2022), and an Ensemble model based on the previous three individual models was also fit (*fit_ensemble, flexsdm)* (Velazco et al. 2022).The same performance measures as for vectors were used to select the best performing model.

Model predictions across Europe for both vectors and hosts were made at a resolution of 0.00833° which equates to about 1km^2^ at the equator. Model predictions were made in R (v. 4.4.1) (flexsdm::*sdm_predict,* v1.3.4, Velazco et al. 2022) using the highest scoring model as selected in Withers et al. (2024).Table 1 A series of risk maps were then produced to estimate WNV risk across Europe based on habitat suitability, with the same process being used for both the current and 2100 climatic and environmental conditions under SSP1 and SSP5, as outlined in Withers et al. (2024) and briefly here. Initially, a vector presence likelihood risk map was produced by multiplying the habitat suitability scores of cells for C*x. pipiens* and *Cx. modestus*. For a host presence likelihood map the suitability scores of all cells overlayed were summed to give an overall host suitability score of both representative corvid and non-corvid groups (probability of either host group being present). These vector and host maps were then combined by multiplying the host and vector maps together to produce an overall WNV risk map (probability of both vectors and hosts being present). Then, to further define risk in terms of human and equine health, WNV risk maps were overlayed separately with human population density for the current and future conditions (Gao 2020, Gao 2017), and current horse population density (Gilbert et al. 2022).

**Table 1:** Model performance scores of the selected vector and avian host models. The performance metrics are True positive rate (TPR), True negative rate (TNR), Sorensen, Jaccard, F-measure (FPB), Omission rate (OR), True skill statistic (TSS), Area under curve (AUC), Boyce index and Inverse mean absolute error (IMAE). Higher values indicate better model performance.

|  |  | TPR |  | TNR |  | Sorensen |  | Jaccard |  | FPB |  | OR |  | TSS |  | AUC |  | Boyce |  | IMAE |  |
| --- | --- | --- | --- | --- | --- | --- | --- | --- | --- | --- | --- | --- | --- | --- | --- | --- | --- | --- | --- | --- | --- |
|  | score range | 0 to 1 |  | 0 to 1 |  | 0 to 1 |  | 0 to 1 |  | 0 to 2 |  | 0 to 1 |  | -1 to 1 |  | 0 to 1 |  | -1 to 1 |  | 0 to 1 |  |
|  |  | mean | sd | mean | sd | mean | sd | mean | sd | mean | sd | mean | Sd | mean | sd | mean | sd | mean | sd | mean | sd |
| Culex pipiens |  |  |  |  |  |  |  |  |  |  |  |  |  |  |  |  |  |  |  |  |  |
| Ensemble mean weighted |  | 0.65 | 0.21 | 0.77 | 0.10 | 0.52 | 0.17 | 0.36 | 0.15 | 0.72 | 0.30 | 0.35 | 0.21 | 0.42 | 0.11 | 0.76 | 0.08 | 0.95 | 0.02 | 0.68 | 0.01 |
| Culex modestus |  |  |  |  |  |  |  |  |  |  |  |  |  |  |  |  |  |  |  |  |  |
| Neural Network |  | 0.79 | 0.17 | 0.61 | 0.16 | 0.53 | 0.03 | 0.36 | 0.03 | 0.72 | 0.06 | 0.21 | 0.17 | 0.39 | 0.01 | 0.68 | 0.08 | 0.93 | 0.08 | 0.67 | 0.02 |
| Corvid |  |  |  |  |  |  |  |  |  |  |  |  |  |  |  |  |  |  |  |  |  |
| GBM |  | 0.77 | 0.05 | 0.82 | 0.04 | 0.79 | 0.02 | 0.65 | 0.03 | 1.31 | 0.07 | 0.23 | 0.05 | 0.59 | 0.08 | 0.86 | 0.04 | 0.99 | 0.01 | 0.70 | 0.02 |
| Non-corvid |  |  |  |  |  |  |  |  |  |  |  |  |  |  |  |  |  |  |  |  |  |
| Ensemble mean weighted |  | 0.85 | 0.03 | 0.84 | 0.02 | 0.85 | 0.04 | 0.74 | 0.06 | 1.47 | 0.11 | 0.15 | 0.03 | 0.70 | 0.04 | 0.92 | 0.01 | 1.00 | 0.00 | 0.77 | 0.01 |

When predicting species distribution models, it is important to consider model transferability to geographic regions outside of the extent of the training data. This is important as the environmental conditions may be different to those the model was built on, meaning the model must extrapolate into novel conditions. Here, model transferability was determined using Mahalanobis distance and the shape method (Velazco et al. 2022, Velazco et al. 2023). In this method, a high Shape value represents areas with a greater environmental distance from the training data so areas with higher values will have less reliable predictions than those with lower Shape values (Velazco et al. 2023).

## Results

### Model selection

The highest scoring models for each vector and host group were selected from Withers et al. (2024). These were the mean-weighted Ensemble model for *Cx. pipiens,* the Neural Network model for *Cx. modestus,* the Generalized Boosted Regression model for corvids and the mean-weighted Ensemble model for non-corvids (Table 1, full details of all models are available in Supplementary Information S1).

### Extrapolation across Europe

The environmental conditions that were covered by the presence data for both *Cx. pipiens* and *Cx. modestus* was generally representative of those found across the European study area, with much of the study area having low extrapolation scores (Figures **Figure 1**). Environmental representation was better for *Cx. pipiens* compared to *Cx. modestus,* however, as more presence data were available for *Cx. pipiens* this was expected. Similar patterns of environmental representation were observed when predicting into 2100 under both SSP1 and SSP5, with the lowest representation for the worst-case climate scenario for both *Cx. pipiens* and *Cx. modestus* (Figures **Figure 1**). Though some areas are less represented for both vector species, a wide range of the environmental conditions in the study area were represented overall, enabling us to process with Europe-wide predictions of suitability.

**Figure 1:**
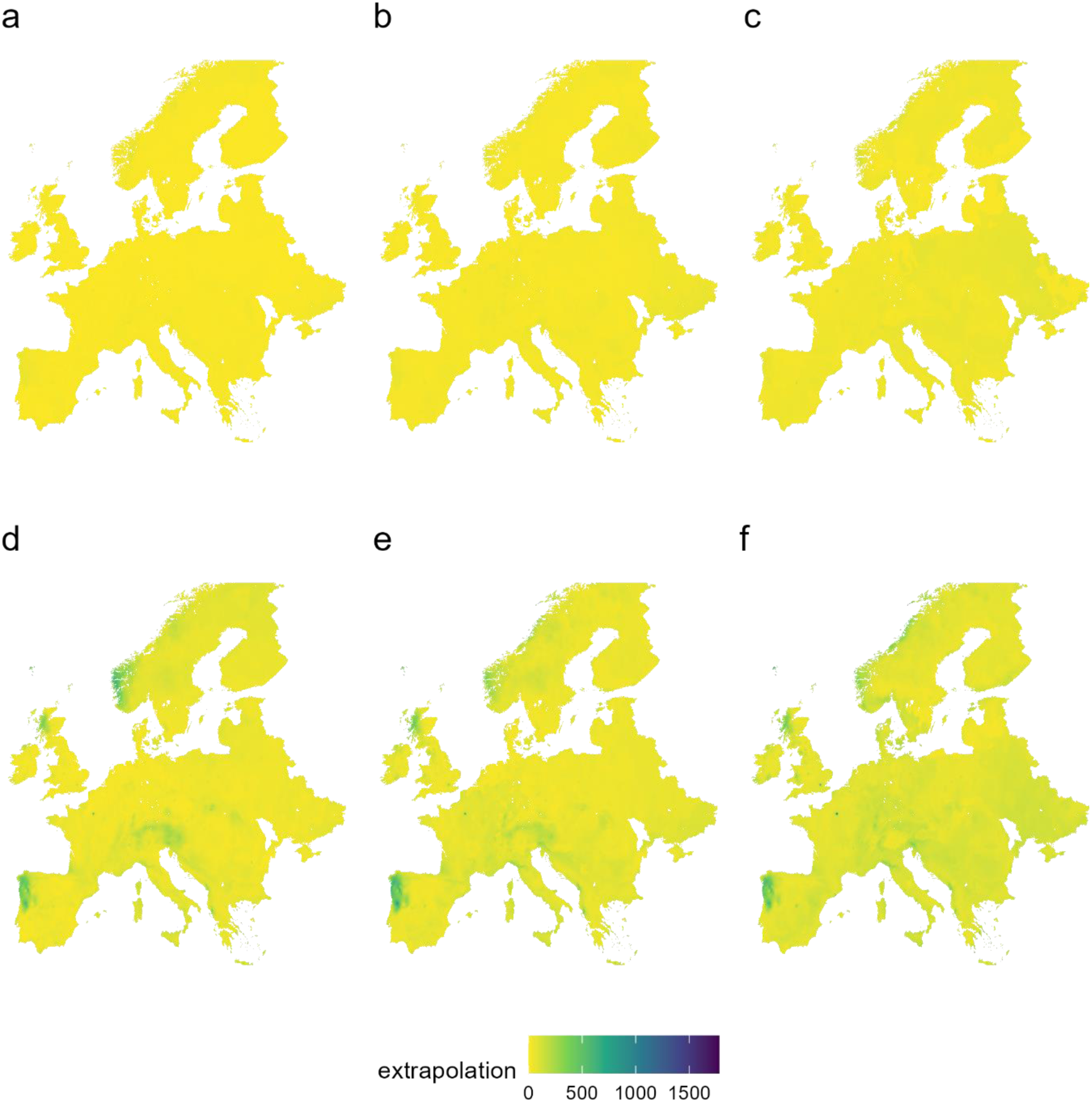
Shape measure of extrapolation for a) Cx. pipiens current, b) Cx. pipiens SSP1 2100, c) Cx. pipiens SSP5 2100, d) Cx. modestus current, e) Cx. modestus SSP1 2100 and f) Cx. modestus SSP5 2100. The higher the shape value the more novel the environmental conditions being predicted, meaning darker colours (i.e., blues and purples) represent areas that were more poorly represented by the data used to train the models.

### Predicting environmental suitability for *Culex* vectors across Europe using correlative species distribution models

Environmental suitability for *Cx. pipiens* was widespread across much of Europe, with hotspots of higher suitability (e.g., southeast England, Belgium, northern Italy) (**Figure 1a**). This pattern of suitability was also projected into 2100 for both best (SSP1) and worst (SSP5) case scenarios, however suitability was higher and more widespread in the future under both conditions, with the greatest suitability (i.e., more widespread, more hotspots) predicted to be under SSP5 (**Figure 1b and 1c**).

Environmental suitability was generally lower and less widespread for *Cx. modestus* compared to *Cx. pipiens* across Europe. Hotspots of suitability were present across Europe, and some of these were the same as for *Cx. pipiens* (e.g., southeast England, Netherlands) however some were different (e.g., southern Spain, southern Italy) highlighting the different environmental niches of these two *Culex* species (**Figure 1d**). Similar patterns of suitability were predicted into 2100 for *Cx. modestus,* again with future predictions showing increased range and higher suitability for both SSP1 and SSP5, with SSP5 leading to greater suitability (**Figure 1e and 1f**).

As risk is likely to be higher where *Cx. pipiens* and *Cx. modestus* are both present, the environmental suitability of these two vectors was combined to produce a single vector risk map for WNV across Europe for the current environment (**Figure 3a**). This showed that generally across Europe there were areas of high suitability for both vectors, and these areas increased in suitability when considering the future environment and climate (**Figure 3b and 3c**).

**Figure 2:**
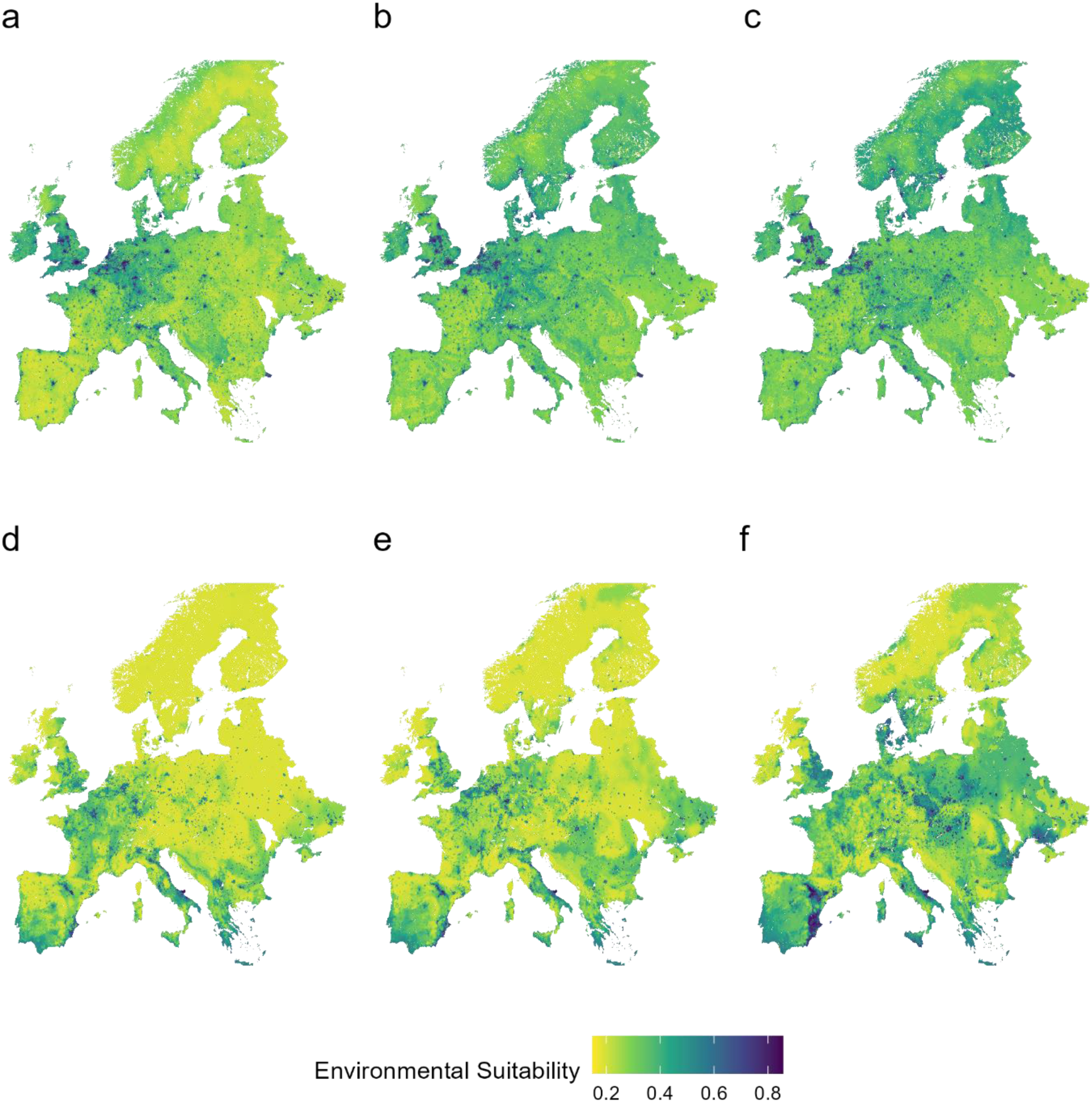
Environmental suitability of Culex pipiens in **a)** the current climate, **b)** 2100 under SSP1 **c)** 2100 under SSP5 and Culex modestus in **d)** the current climate, **e)** 2100 under SSP1 and **f)** 2100 under SSP5.

**Figure 3:**
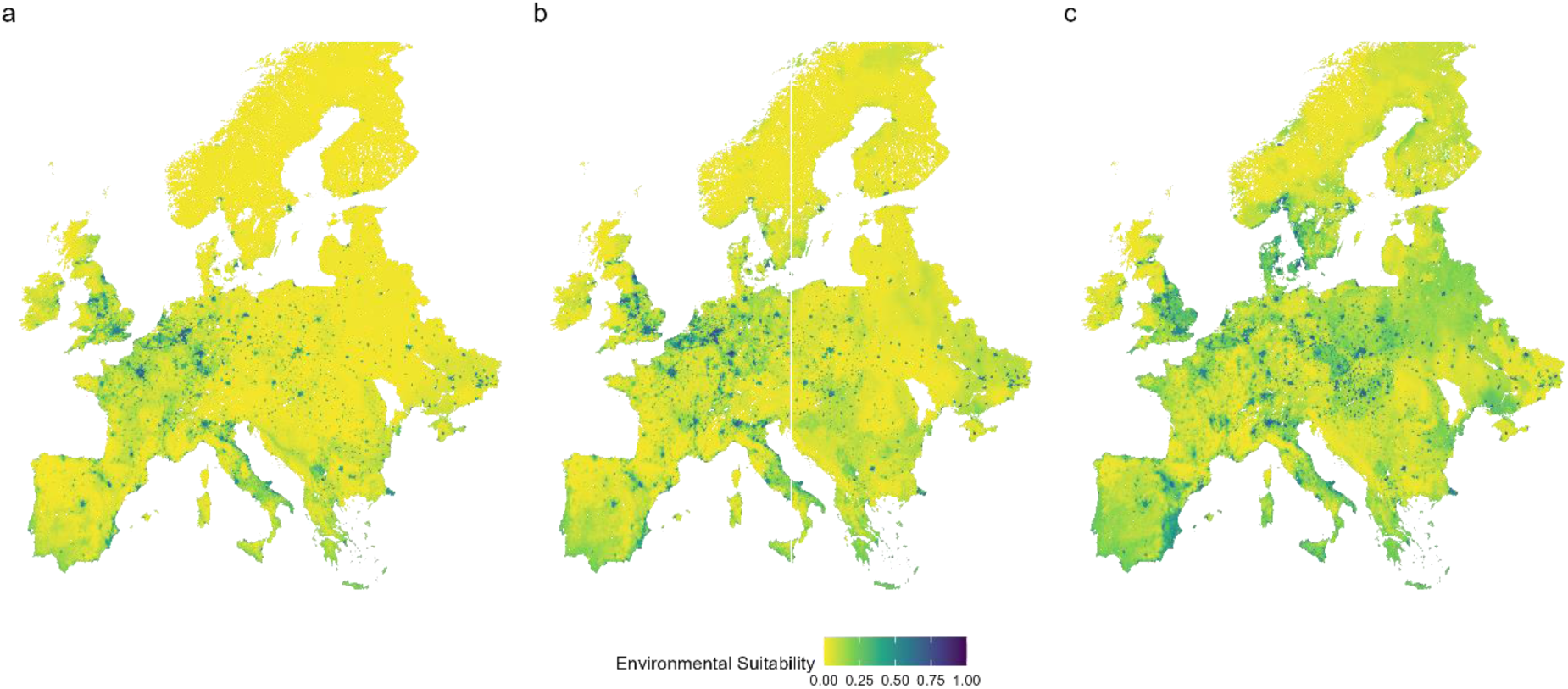
Environmental suitability of Europe for both Cx. pipiens and Cx. modestus combined, for the a) the current climate, b) 2100 under SSP1 and c) 2100 under SSP5.

### Predicting environmental suitability for *Culex* vectors across Europe using a mechanistic species distribution model, CLIMEX

Suitability for both *Cx. pipiens* and *Cx. modestus* was predominantly driven by temperature in the CLIMEX model, with areas of high elevation in Scotland, Scandinavia and across the alps all showing very low suitability for both species as an ecoclimatic index score of 0 would suggest no population growth could occur (**Figure 4a-b**). Areas of higher suitability were typically coastal regions of southern and western mainland Europe, with much of the rest of Europe showing as suitable for population growth for both species, with *Cx. pipiens* having a wider range than *Cx. modestus* (**Figure 4a-b**).

**Figure 4:**
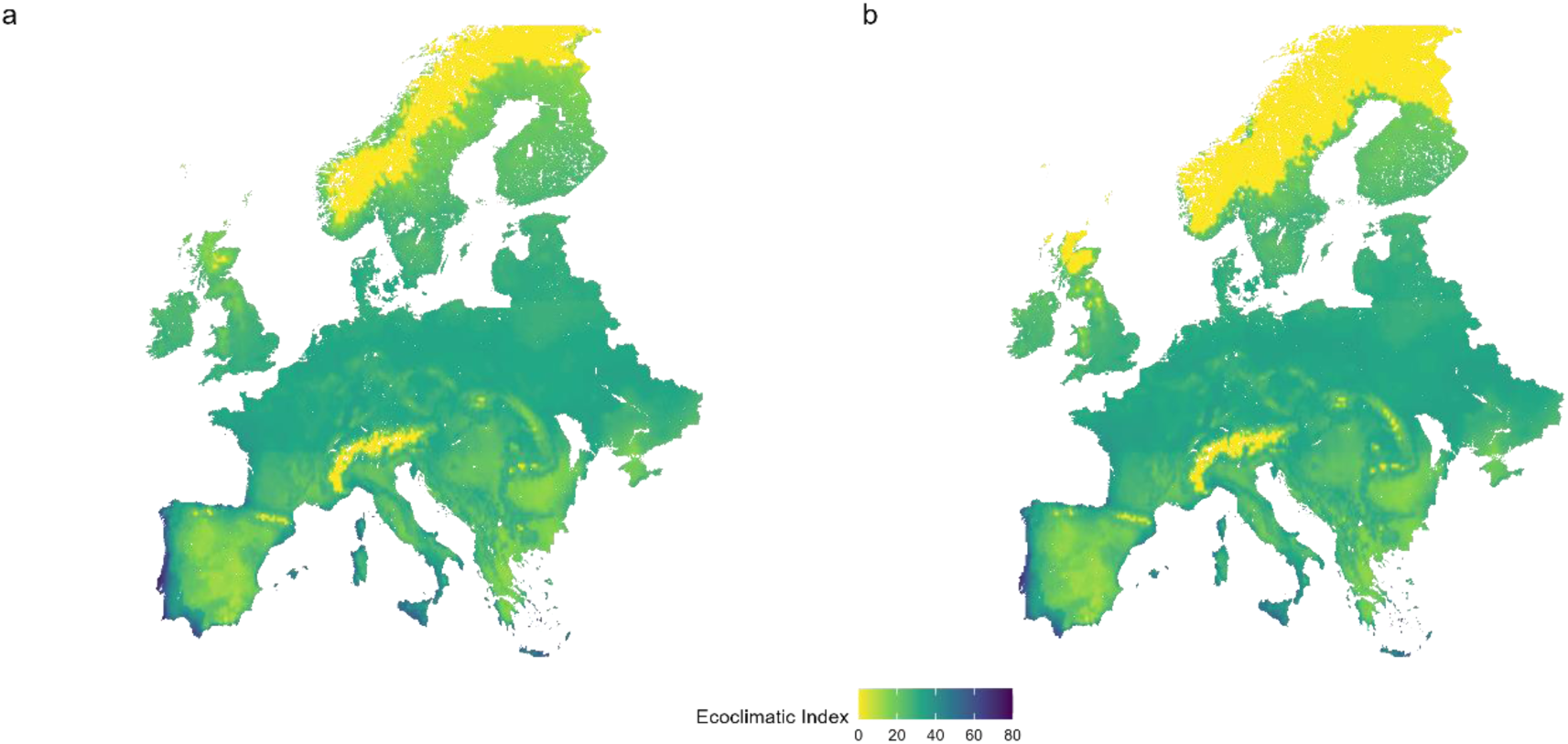
The ecoclimatic index for a) Cx. pipiens and b) Cx. modestus. An ecoclimatic index greater than 1 indicates some survival can occur, with the higher the number the more suitable the area is climatically for the species.

Validation to known presence locations based on ecoclimatic index scores for *Cx. pipiens* showed that the CLIMEX model was largely accurate, with 911 presence points falling within “very favourable areas”, 20 in “favourable areas”, and just 5 falling into “marginally suitable areas” or “unsuitable areas”. Similarly, for *Cx. modestus,* all 44 available presence points within the study area fell within areas considered to be “very favourable”.

### Predicting environmental suitability for avian hosts across Europe

There was very high suitability for hosts across western Europe, with some lower suitability predicted for southern Spain and across much of Eastern Europe including Finland, Poland and Ukraine (**Figure 5**). When predicting into 2100 for both SSP1 and SSP5 there were some areas that saw an increase in suitability (e.g., southern Finland, Estonia, Switzerland), however other areas saw a decrease in suitability (e.g., eastern Spain, France) (**Figure 5b and 5c**).

**Figure 5:**
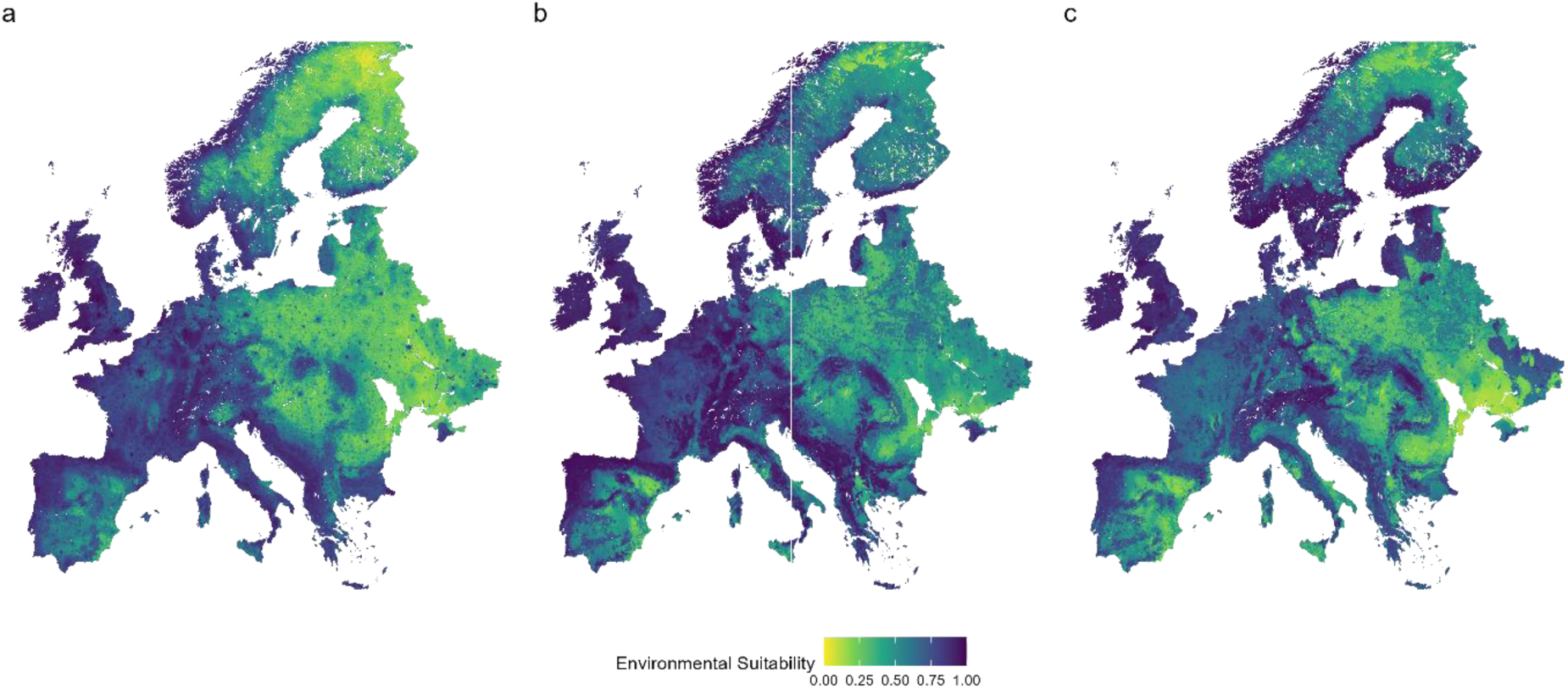
Environmental suitability of Europe for both corvid and non-corvid representative avian host groups combined, for the **a)** the current climate, **b)** 2100 under SSP1 and **c)** 2100 under SSP5.

### WNV risk considering vector and host suitability

The greatest risk of WNV will be in areas where environmental suitability is high for both vectors and hosts. This means there are areas of much higher risk across Europe, and these areas are generally located around more urban or wetland areas which are favoured by the vectors (**Figure 6a**). When predicting into the future, some areas see an increase in suitability for both vectors and hosts leading to higher risk, such as central England, northern Belgium and west Sweden (**Figure 6b-d**). However, other areas see a decline in environmental suitability, and this leads to reduced WNV risk, for example coastal regions in south-east England, northern Italy and western Germany (**Figure 6b-d**).

**Figure 6:**
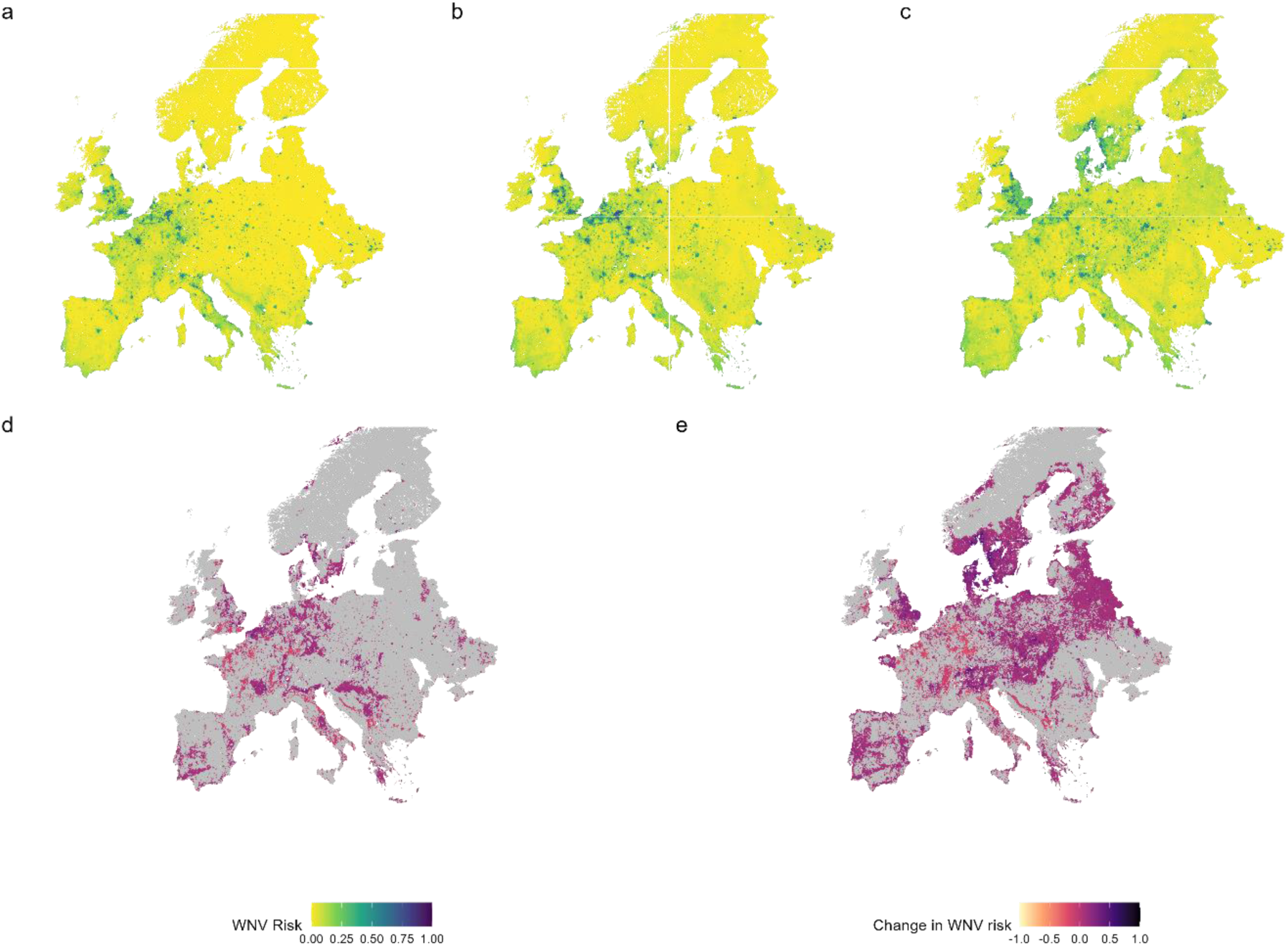
WNV risk considering vector and avian host suitability for **a)** the current environmental conditions, **b)** 2100 under SPP1, **c)** 2100 under SSP5, and the change in risk by 2100 under **d)** SPP1 and **e)** SSP5. Grey areas on c and d represent areas with very little or no change (-0.05 to 0.05).

### Determining WNV risk for horses and humans

The risk of WNV was considered in relation to human and horse density, as these dead-end hosts are at greatest risk in terms of impacts on human and animal health and the economy. As expected, the greatest risk to horses coincides with areas of higher suitability for both avian hosts and vectors, with patches across England, Belgium and Germany (**Figure 7**). When considering the risk to human health, the greatest risk across Europe is predicted to be in urbanised areas that also have higher suitability for vectors and hosts (**Figure 8a**). When predicting risk in 2100 under both SSP1 and SSP5, the general areas at risk remain similar, such as London in England and Paris in France, as they are predominantly driven by urbanised areas of higher human population density that are also highly suitable for vectors and hosts (**Figure 8b-c**).

**Figure 7:**
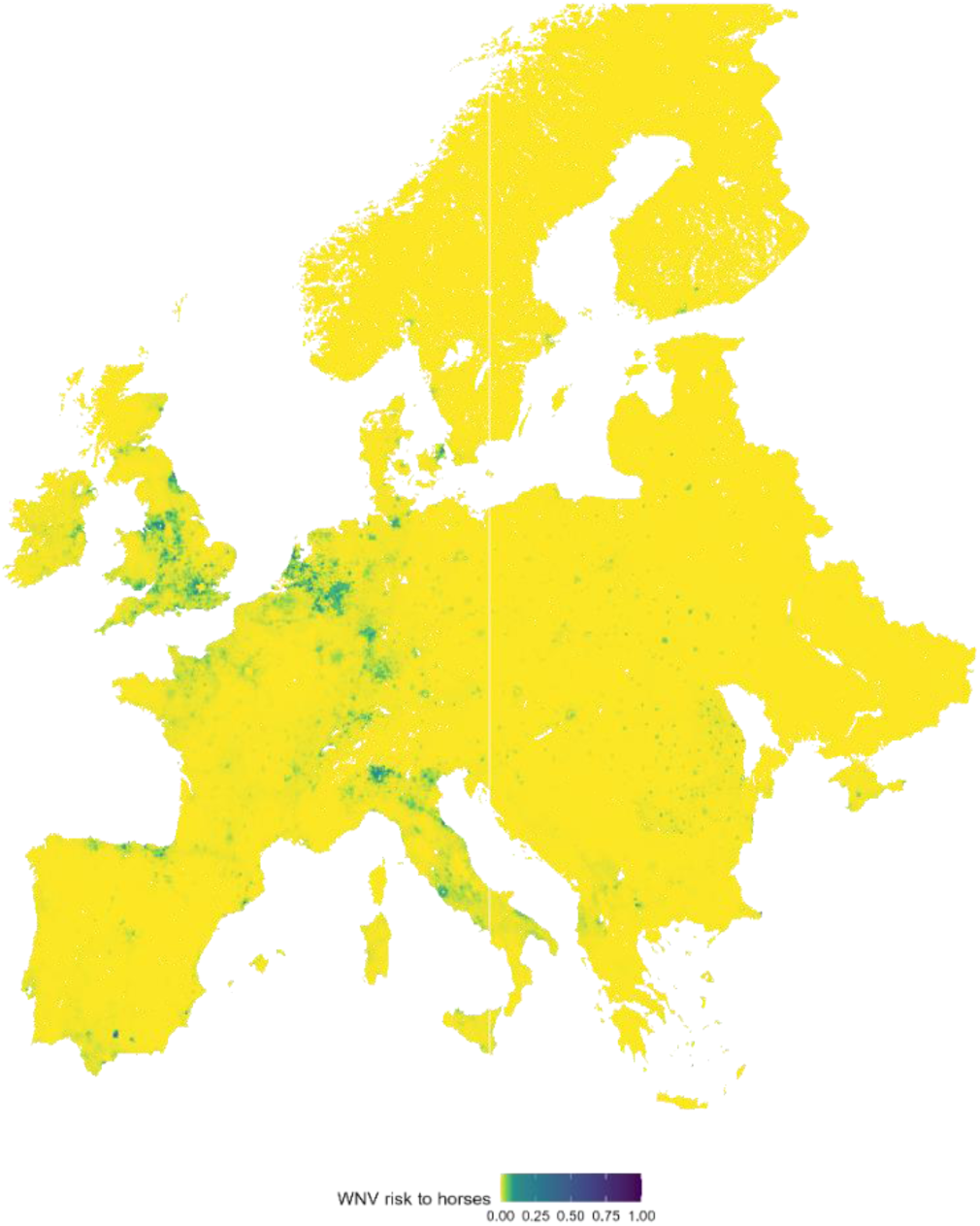
The WNV risk in Europe calculated by considering vector (Cx. pipiens, Cx. modestus), target reservoir host (corvid and non-corvid) and the risk to horses in the current climate.

**Figure 8:**
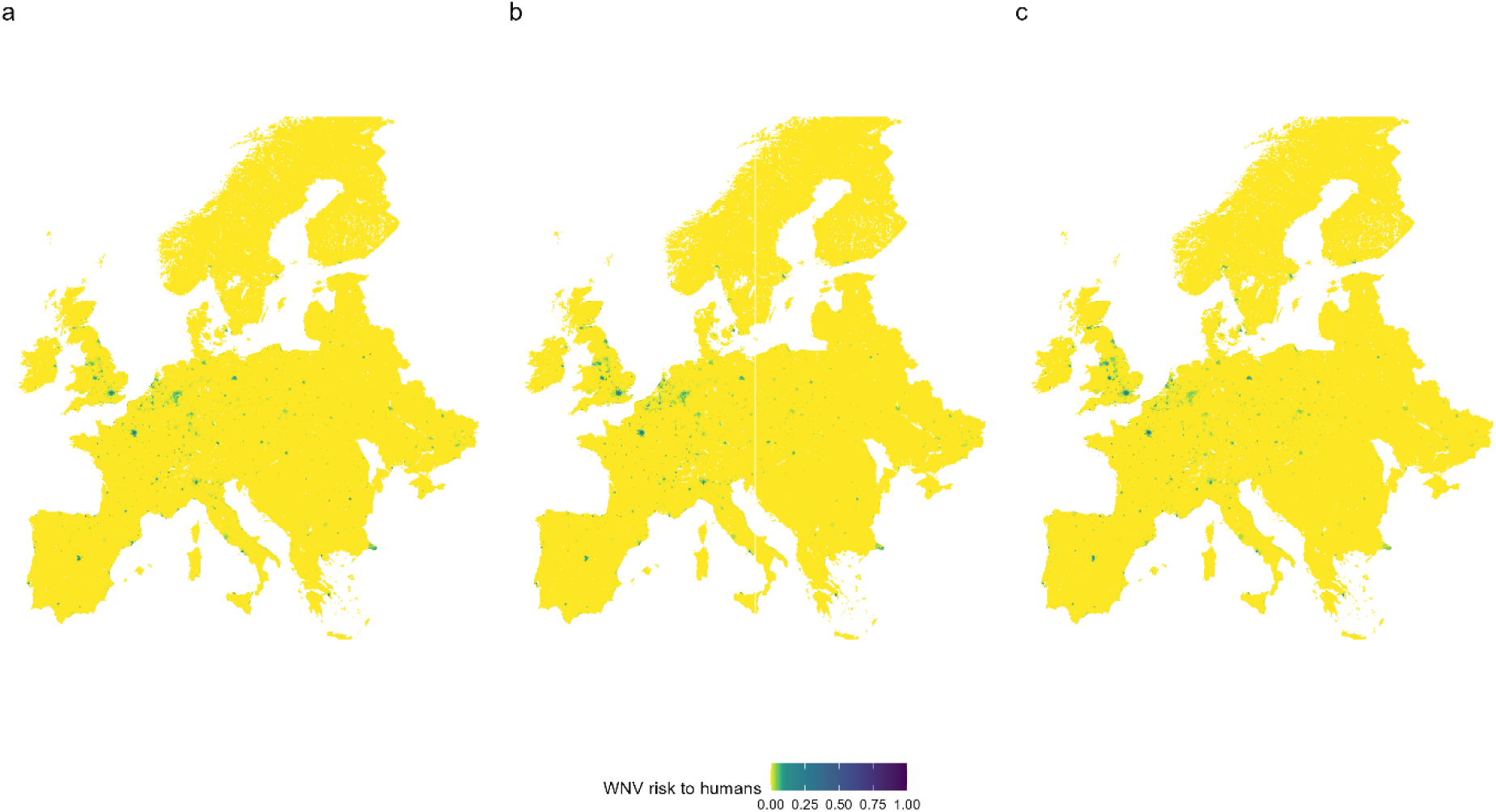
The WNV risk in Europe calculated by considering vector (Cx. pipiens, Cx. modestus), target reservoir host (corvid and non-corvid) and human population density. The figure shows a) the risk to the human population in the current climate, b) the risk to the human population in 2100 under scenario SSP1 and, c) the risk to the human population in 2100 under scenario SSP5.

## Discussion

This study has demonstrated that species distribution modelling can be used to predict areas at higher risk of mosquito-borne diseases across Europe, using correlative species distribution models of arthropod vectors that have been validated by mechanistic species distribution models. These vector models have then been combined with species distribution models for avian hosts, and density estimates of human and equines to give an overall estimate of risk of infection to dead-end hosts.

Due to the presence of WNV in parts of mainland Europe included in this study area, visual comparisons can also be made to where locally acquired human WNV infections have been reported in 2024 (European Centre for Disease Prevention and Control 2024). Some regions where WNV has been reported are not predicted to be at high risk in the approach here, such as Eastern Europe (e.g., Ukraine, Belarus), however, these areas have less data available for model training meaning that extrapolation is less reliable. Overall, in other areas both risk maps presented here, and the human incidence maps align. For example, regions throughout Italy and northern Germany are predicted to be at greater risk and have had WNV cases reported (European Centre for Disease Prevention and Control 2024). These alignments between locally acquired human incidence cases and areas predicted to be at higher risk further validate these disease risk models and demonstrate that the approach outlined in this work, and previous work by Withers et al. (2024), can be useful in highlighting areas at greater risk of novel vector-borne disease.

Across Europe, there are patches of higher and lower risk that are predominantly driven by vector suitability as avian hosts are widespread. This is expected as the mosquito vectors have more niche environmental requirements, including temperature, precipitation and land use meaning the species distribution models show less widespread suitability than the models for the corvid and non-corvid representative hosts (Ciota et al., 2014, Ewing et al. 2017, Golding et al. 2012, Golding et al. 2015, Jones et al. 2012). This also leads to hotspots of higher risk for both human and equine health, for example, for the human WNV risk maps the highest risk is predicted to be localised to urbanised areas of higher population density, particularly in areas with fresh water such as London in southern England and Paris in northern France. It should be noted that whilst these maps can predict environmental change, they are unable to consider human behaviour which may increase the risk further, such as an increase in water storage tanks that has been proposed to increase mosquito (*Aedes aegypti*) populations and consequently increase disease risk (Beebe et al. 2009).

Considering the future the general pattern of areas of higher and lower risk remains unchanged, however across Europe some areas see an increase in risk whilst some areas see a decrease in risk. This suggests that the overall impact of environmental change will not be consistent across the continent, as changes will see some areas become less suitable for vectors and hosts and other areas become more suitable. For example, northern Europe is predicted to see an increase in suitability for both vectors and hosts projecting into the future under both SSP1 and SSP5 scenarios which is likely driven in part by a warming climate, and consequently an increase in disease risk is forecast. Similar changes in risk have also been predicted for other mosquito species, such as *Anopheles gambiae* in Africa, with climate change decreasing suitability in western Africa whilst increasing suitability in eastern and southern Africa (Peterson 2009).

Mechanistic models produced using CLIMEX are a valuable tool in predicting vector distributions (Early et al. 2022, Khormi & Kumar 2014, Kriticos et al. 2015, Poutsma et al. 2008, Sutherst & Maywald 1985). However, for some vectors variables that cannot be included in mechanistic models are important, such as human density and land-use (Ferraguti et al. 2023). Therefore, here we used mechanistic models to validate that areas predicted to have higher suitability in correlative models were within the biological limits of *Culex* vectors. We found that those areas predicted to be of lower suitability in correlative models also had low ecoclimatic index scores in the mechanistic models, and similarly areas predicted to be of high suitability in correlative models had higher ecoclimatic index scores.

Both modelling approaches presented here would be improved by further studies to collect more data on *Culex* abundance and biological limitations. Mechanistic modelling relies on detailed information on a species’ survival limitations (e.g., upper and lower temperature limits), and whilst this was relatively well covered for *Cx. pipiens* (Ewing et al. 2016, Field et al. 2022, Ewing et al. 2019), there are many gaps in our knowledge for *Cx. modestus* (Soto & Delang 2023). These knowledge gaps made mechanistic modelling a challenge with more parameters based on similar species (i.e., *Cx. pipiens*) so the outputs are likely to be less representative of the true potential range. Correlative modelling largely relies on presence-absence data and false absences can greatly influence the models (Croft & Smith 2019, Liu et al. 2019). However, as is the case for many species, for both *Cx. pipiens* and *Cx. modestus* there was insufficient absence data available meaning that pseudo-absence data had to be used instead, and it was decided here to use randomly selected background points as this had the fewest assumptions about the species (Valavi et al. 2021, Valavi et al. 2022, Elith et al. 2006, VanDerWal et al. 2009).

Overall, this study has shown that the modelling approach developed for the UK can be generalised for use over a wider area to predict areas of disease risk for WNV. By using species distribution models of vectors and both target and non-target hosts a clearer picture of risk can be developed as areas of high suitability for both will have higher risk. Nevertheless, other factors may influence risk of emergence such as the existence of introduction pathways and virus emergence in adjacent areas. Future work can expand this approach to look at novel diseases and vectors, both on a more localised and wider scale.

## Supplementary Information (S1)

Several different performance measures were used to determine the most suitable model for each vector and host group (Table S1). The performance metrics used were True positive rate (TPR), True negative rate (TNR), Sorensen, Jaccard, F-measure of presence-background (FPB), Omission rate (OR), True skill statistic (TSS), Area under curve (AUC), Boyce index and Inverse mean absolute error (IMAE) (Velazco et al., 2022, Valavi et al., 2022, Valavi et al., 2021, Hijmans and Elith, 2013, Roberts et al., 2017).

**Table S1:**
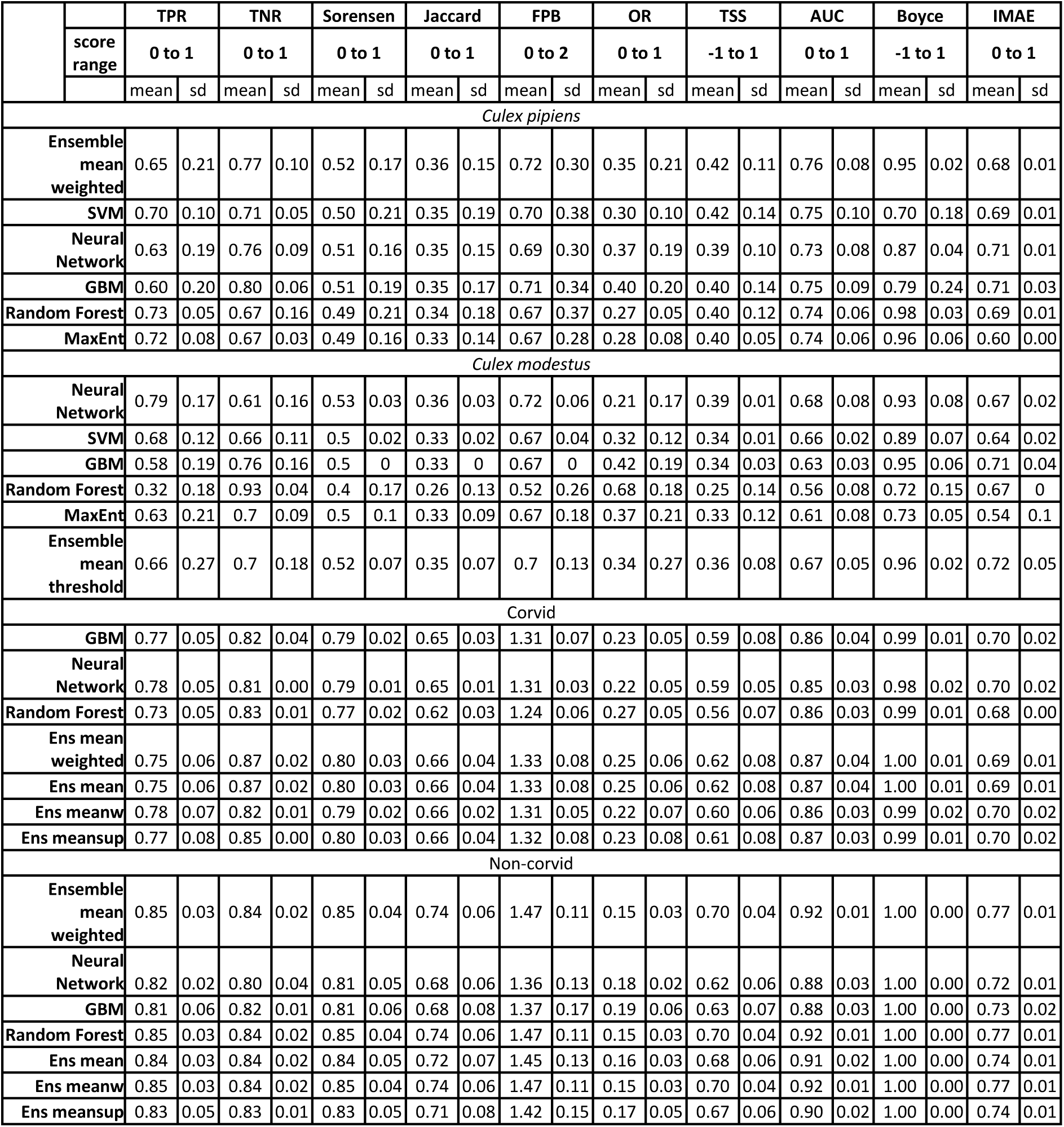
Model performance scores for all the vector and avian host models. The performance metrics are True positive rate (TPR), True negative rate (TNR), Sorensen, Jaccard, F-measure (FPB), Omission rate (OR), True skill statistic (TSS), Area under curve (AUC), Boyce index and Inverse mean absolute error (IMAE). Higher values indicate better model performance.

## References

Amdouni J, Conte A, Ippoliti C, Candeloro L, Tora S, Sghaier S, Hassine TB, Fakhfekh EA, Savini G, Hammami S (2022). Culex pipiens distribution in Tunisia: Identification of suitable areas through Random Forest and MaxEnt approaches. Veterinary Medicine and Science, 8, 2703–2715.

Baici JE, Bowman J (2023). Combining community science and MaxEnt modeling to estimate Wild Turkey (*Meleagris gallopavo*) winter abundance and distribution. Avian Conservation and Ecology, 18.

Beebe NW, Cooper RD, Mottram P, Sweeney AW (2009). Australia’s Dengue risk driven by human adaptation to climate change. PLOS Neglected Tropical Diseases, 3, e429.

Bessell PR, Robinson RA, Golding N, Searle KR, Handel IG, Boden LA, Purse BV, Bronsvoort BMDC (2016). Quantifying the risk of introduction of West Nile Virus into Great Britain by migrating Passerine birds. Transboundary and Emerging Diseases, 63, 347–359.

Chapman D, Pescott OL, Roy HE, Tanner R (2019). Improving species distribution models for invasive non-native species with biologically informed pseudo-absence selection. Journal of Biogeography, 46, 1029–1040.

Ciota AT, Matacchiero AC, Kilpatrick AM, Kramer LD (2014). The effect of temperature on life history traits of Culex mosquitoes. Journal of Medical Entomology, 51, 55–62.

Constant O, Gil P, Barthelemy J, Bolloré K, Foulongne V, Desmetz C, Leblond A, Desjardins I, Pradier S, Joulié A, Sandoz A, Amaral R, Boisseau M, Rakotoarivony I, Baldet T, Marie A, Frances B, Reboul Salze F, Tinto B, Van De Perre P, Salinas S, Beck C, Lecollinet S, Gutierrez S, Simonin Y (2022). One Health surveillance of West Nile and Usutu viruses: a repeated cross-sectional study exploring seroprevalence and endemicity in Southern France, 2016 to 2020. Eurosurveillance, 27, 2200068.

Croft S, Chauvenet ALM, Smith GC (2017). A systematic approach to estimate the distribution and total abundance of British mammals. PLOS ONE, 12, e0176339.

Croft S, Smith GC (2019). Structuring the unstructured: estimating species-specific absence from multi-species presence data to inform pseudo-absence selection in species distribution models. bioRxiv, 656629.

Culverwell CL, Vapalahti O (2023). First record of *Culex modestus* in Finland. Journal of the European Mosquito Control Association, 41, 63–66.

Early R, Rwomushana I, Chipabika G, Day R (2022). Comparing, evaluating and combining statistical species distribution models and CLIMEX to forecast the distributions of emerging crop pests. Pest Management Science, 78, 671–683.

Elith J, Graham CH, Anderson RP, Dudík M, Ferrier S, Guisan A, Hijmans RJ, Huettmann F, Leathwick, JR, Lehmann A, Li J, Lohmann LG, Loiselle BA, Manion G, Moritz C, Nakamura M, Nakazawa Y, Overton JM, Townsend Peterson A, Phillips SJ, Richardson K, Scachetti-Pereira R, Schapire RE, Soberón J, Williams S, Wisz MS, Zimmermann NE (2006). Novel methods improve prediction of species’ distributions from occurrence data. Ecography, 29, 129–151.

Elith J, Kearney M, Phillips S (2010). The art of modelling range-shifting species. Methods in Ecology and Evolution, 1, 330–342.

Endo A, Amarasekare P (2022). Predicting the spread of vector-borne diseases in a warming world. Frontiers in Ecology and Evolution, 10.

European Centre for Disease Prevention and Control (2019). A spatial modelling method for vector surveillance.

European Centre for Disease Prevention and Control. (2024). Surveillance of West Nile virus infections in humans, weekly report [Online]. ECDC. Available: https://www.ecdc.europa.eu/en/west-nile-fever/surveillance-and-disease-data/disease-data-ecdc [Accessed 18/12/2024].

Ewing DA, Cobbold CA, Purse BV, Nunn MA, White SM (2016). Modelling the effect of temperature on the seasonal population dynamics of temperate mosquitoes. Journal of Theoretical Biology, 400, 65–79.

Ewing DA, Purse BV, Cobbold CA, Schäfer SM, White SM (2017). Seasonal abundance data of all *Culex pipiens* life stages at a UK field site.

Ewing DA, Purse BV, Cobbold CA, Schäfer SM, White SM (2019). Uncovering mechanisms behind mosquito seasonality by integrating mathematical models and daily empirical population data: *Culex pipiens* in the UK. Parasites & Vectors, 12, 74.

Faburay B 2015. The case for a “one health” approach to combating vector-borne diseases. Infection Ecology & Epidemiology, 5, 28132.

Ferraguti M, Magallanes S, Suarez-Rubio M, Bates PJJ, Marzal A, Renner SC (2023). Does land-use and land cover affect vector-borne diseases? A systematic review and meta-analysis. Landscape Ecology, 38, 2433–2451.

Field EN, Shepard JJ, Clifton ME, Price KJ, Witmier BJ, Johnson K, Boze B, Abadam C, Ebel GD, Armstrong PM, Barker CM, Smith RC (2022). Semi-field and surveillance data define the natural diapause timeline for Culex pipiens across the United States. Communications Biology, 5, 1300.

Furlong M, Adamu A, Hickson RI, Horwood P, Golchin M, Hoskins A, Russell T (2022). Estimating the distribution of Japanese Encephalitis vectors in Australia using ecological niche modelling. Tropical Medicine and Infectious Disease, 7.

Gao J (2017). Downscaling global spatial population projections from 1/8-degree to 1-km grid cells. National Center for Atmospheric Research, Boulder, CO, USA.

Gao J (2020). Global 1-km downscaled population base year and projection grids based on the shared socioeconomic pathways, Revision 01. NASA Socioeconomic Data and Applications Center (SEDAC).

GBIF (2022). Occurrence Download: Culex. 10.15468/dl.fqc9t3

GBIF (2024). Occurrence Download - Turdus merula. 10.15468/DL.FVVQFT

GBIF (2024a). Occurrence Download - Corvus corone. 10.15468/DL.Q7TAWD

GBIF (2024b). Occurrence Download - Turdus merula. 10.15468/DL.7MYK3H

GBIF (2024c). Occurrence Download - Corvus corax. 10.15468/DL.QZ6KFF

Gilbert M, Cinardi G, Da Re D, Wint WGR, Wisser D, Robinson TP (2022). Global horse distribution in 2015 (5 minutes of arc).

Golding N, Nunn MA, Medlock JM, Purse BV, Vaux AGC, Schäfer SM (2012). West Nile virus vector *Culex modestus* established in southern England. Parasites & Vectors, 5, 32.

Golding N, Nunn MA, Purse BV (2015). Identifying biotic interactions which drive the spatial distribution of a mosquito community. Parasites & Vectors, 8, 367.

Gorris ME, Bartlow AW, Temple SD, Romero-Alvarez D, Shutt DP, Fair JM, Kaufeld KA, Del Valle SY, Manore CA (2021). Updated distribution maps of predominant *Culex* mosquitoes across the Americas. Parasites & Vectors, 14, 547.

Hijmans RJ, Elith J (2013). Species Distribution Modeling with R, Elsevier Inc.

Hijmans RJ, Phillips S, Leathwick J, Elith J (2023). Dismo: Species Distribution Modeling. 1.3–14 ed.

Humblet MF, Vandeputte S, Fecher-Bourgeois F, Léonard P, Gosset C, Balenghien T, Durand B, Saegerman C (2016). Estimating the economic impact of a possible equine and human epidemic of West Nile virus infection in Belgium. Eurosurveillance, 21(31), 30309.

Jones CE, Lounibos LP, Marra PP, Kilpatrick AM (2012). Rainfall influences survival of *Culex pipiens* in a residential neighborhood in the mid-Atlantic United States. Journal of Medical Entomology, 49, 467–473.

Khormi HM, Kumar L (2014). Climate change and the potential global distribution of *Aedes aegypti*: spatial modelling using geographical information system and CLIMEX. Geospatial Health, 8, 405–415.

Koch FH (2021). Considerations regarding species distribution models for forest insects. Agricultural and Forest Entomology, 23, 393–399.

Kraemer MUG, Reiner R C, Brady OJ, Messina JP, Gilbert M, Pigott DM, Yi D, Johnson K, Earl L, Marczak LB, Shirude S, Davis Weaver N, Bisanzio D, Perkins TA, Lai S, Lu X, Jones P, Coelho GE, Carvalho RG, Van Bortel W, Marsboom C, Hendrickx G, Schaffner F, Moore CG, Nax HH, Bengtsson L, Wetter E, Tatem AJ, Brownstein JS, Smith DL, Lambrechts L, Cauchemez S, Linard C, Faria NR, Pybus OG, Scott TW, Liu Q, Yu H, Wint GRW, Hay SI, Golding N (2019). Past and future spread of the arbovirus vectors *Aedes aegypti* and *Aedes albopictus*. Nature Microbiology, 4, 854–863.

Kriticos DJ, Maywald GF, Yonow T, Zurcher EJ, Herrmann NI, Sutherst RW (2015). CLIMEX Version 4: Exploring the effects of climate on plants, animals and diseases. CSIRO, 184.

Liu C, Newell G, White M (2019). The effect of sample size on the accuracy of species distribution models: considering both presences and pseudo-absences or background sites. Ecography, 42, 535–548.

NBN Atlas (2022). Occurance Records -Culicidae [Online]. Available: https://records.nbnatlas.org/occurrences/search?q=lsid:NBNSYS0000040182&fq=occurrence_status:present&nbn_loading=true#tab_mapView [Accessed 07/11/2022 2022].

Noll M, Wall R, Makepeace BL, Vineer HR (2023). Distribution of ticks in the Western Palearctic: an updated systematic review (2015-2021). Parasites & Vectors, 16, 141.

Oliveira S, Capinha C, Rocha J (2023). Predicting the time of arrival of the Tiger mosquito (*Aedes albopictus*) to new countries based on trade patterns of tyres and plants. Journal of Applied Ecology, 60(11), 2362–2374

Ostlund EN, Andresen JE, Andresen M (2000). West Nile Encephalitis. Veterinary Clinics of North America: Equine Practice, 16, 427–441.

Peterson AT (2009). Shifting suitability for malaria vectors across Africa with warming climates. BMC Infectious Diseases, 9, 59.

Poutsma J, Loomans AJM, Aukema B, Heijerman T (2008). Predicting the potential geographical distribution of the harlequin ladybird, Harmonia axyridis, using the CLIMEX model. BioControl, 53, 103–125.

Pradier S, Lecollinet S, Leblond A (2012). West Nile virus epidemiology and factors triggering change in its distribution in Europe. Revue scientifique et technique (International Office of Epizootics), 31, 829–844.

Richman R, Diallo D, Diallo M, Sall AA, Faye O, Diagne CT, Dia I, Weaver SC, Hanley KA, Buenemann M (2018). Ecological niche modeling of *Aedes* mosquito vectors of chikungunya virus in southeastern Senegal. Parasites & Vectors, 11, 255.

Rocklöv J, Dubrow R (2020). Climate change: an enduring challenge for vector-borne disease prevention and control. Nature Immunology, 21, 479–483.

Simons RRL., Croft S, Rees E, Tearne O, Arnold ME, Johnson N (2019). Using species distribution models to predict potential hot spots for Rift Valley Fever establishment in the United Kingdom. PLOS ONE, 14, e0225250.

Soto A, Delang L (2023). *Culex modestus*: the overlooked mosquito vector. Parasites & Vectors, 16, 373.

Sutherst RW, Maywald GF (1985). A computerised system for matching climates in ecology. Agriculture, Ecosystems & Environment, 13, 281–299.

Valavi R, Elith J, Lahoz-Monfort JJ, Guillera-Arroita G (2021). Modelling species presence-only data with random forests. Ecography, 44, 1731–1742.

Valavi R, Guillera-Arroita G, Lahoz-Monfort, JJ, Elith J (2022). Predictive performance of presence-only species distribution models: a benchmark study with reproducible code. Ecological Monographs, 92, e01486.

Vanderwal J, Shoo LP, Graham C, Williams SE (2009). Selecting pseudo-absence data for presence-only distribution modeling: How far should you stray from what you know? Ecological Modelling, 220, 589–594.

Velazco SJE, Rose MB, De Andrade AFA, Minoli I, Franklin J (2022). flexsdm: An r package for supporting a comprehensive and flexible species distribution modelling workflow. Methods in Ecology and Evolution, 13, 1661–1669.

Velazco SJE, Rose MB, De Marco Jr P, Regan HM, Franklin J (2023). How far can I extrapolate my species distribution model? Exploring shape, a novel method. *Ecography*, e06992.

Velu RM, Kwenda G, Bosomprah S, Chisola MN, Simunyandi M, Chisenga CC, Bumbangi FN, Sande NC, Simubali L, Mburu MM, Tembo J, Bates M, Simuunza MC, Chilengi R, Orba Y, Sawa H, Simulundu E (2023). Ecological niche modelling of *Aedes* and *Culex* mosquitoes: a risk map for chikungunya and West Nile viruses in Zambia. Viruses, 15.

Vogels CBF, Hartemink N, Koenraadt CJM (2017). Modelling West Nile virus transmission risk in Europe: effect of temperature and mosquito biotypes on the basic reproduction number. Scientific Reports, 7, 5022.

Withers AJ, Croft S, Budgey R, Warren D, Johnson N (2024). Using correlative and mechanistic species distribution models to predict vector-borne disease risk for the current and future environmental and climatic change: a case study of West Nile virus in the UK. bioRxiv, 2024.09.12.612656.

World Health Organisation (2020). Vector-borne diseases [Online]. Available: https://www.who.int/news-room/fact-sheets/detail/vector-borne-diseases#:~:text=Vector%2Dborne%20diseases%20account%20for,infection%20transmitted%20by%20Anopheline%20mosquitoes. [Accessed 08/03/2024].

